# An adaptive algorithm for fast and reliable online saccade detection

**DOI:** 10.1101/693309

**Authors:** Richard Schweitzer, Martin Rolfs

**Author notes:** Correspondence concerning this article should be addressed to Richard Schweitzer, Department of Psychology, Humboldt-Universität zu Berlin, Rudower Chaussee 18, 12489 Berlin, Germany.

## Abstract

To investigate visual perception around the time of eye movements, vision scientists manipulate stimuli contingent upon the onset of a saccade. For these experimental paradigms, timing is especially crucial, as saccade offset imposes a deadline on the display change. Although efficient online saccade detection can greatly improve timing, most algorithms rely on spatial-boundary techniques or absolute-velocity thresholds, which both suffer from their respective weaknesses: late detections and false alarms. We propose an adaptive, velocity-based algorithm for online saccade detection that surpasses both standard techniques in speed and accuracy and allows the user to freely define detection criteria. Inspired by the Engbert-Kliegl-algorithm for microsaccade detection, our algorithm computes two-dimensional velocity thresholds from variance in preceding fixation samples, while compensating for noisy or missing data samples. An optional direction criterion limits detection to the instructed saccade direction, further increasing robustness. We validated the algorithm by simulating its performance on a large saccade dataset and found that high detection accuracy (false-alarm rates of <1%) could be achieved with detection latencies of only 3 milliseconds. High accuracy was maintained even under simulated high-noise conditions. To demonstrate that purely intra-saccadic presentations are technically feasible, we devised an experimental test, in which a Gabor patch drifted at saccadic peak velocities. While this stimulus was invisible when presented during fixation, observers reliably detected it during saccades. Photodiode measurements verified that – including all system delays – stimuli were physically displayed on average 20 ms after saccade onset. Thus, the proposed algorithm provides valuable tool for gaze-contingent paradigms.

## Introduction

In the field of active vision most eye tracking experiments study visual perception around goal-directed rapid eye movements, so-called saccades. Specifically when investigating trans-saccadic or intra-saccadic perception, an experimental paradigm has to be implemented in a way that a stimulus or configuration of stimuli is manipulated online (i.e., in real-time) and gaze-contingently starting with the onset of a saccade (Higgins & Rayner, 2015; Hollingworth, Richard, & Luck, 2008; Melcher & Colby, 2008; Prime, Vesia, & Crawford, 2011; Wolf & Schütz, 2015). As saccades are rapid and brief events often with a skewed velocity profile (Figure 2a-b), this is not always as trivial as it may initially sound. Every computational step between an eye movement and a change in the display adds undesired delays, and every shortcut (e.g., through rough approximations) may lead to false alarms, that is, the detection of a saccade when none happened. Here we will discuss an algorithm that realizes early online saccade detection without sacrificing reliability and is thus able to greatly reduce the overall delay between saccade onset and display change.

To elucidate the challenge that gaze-contingent paradigms pose with regard to timing, let us consider a typical trans-saccadic experimental scenario: Participants are instructed to make a saccade towards a colored patch at 10 degrees of visual angle (dva) eccentricity, resulting in saccades with average peak velocities of 300 dva per second (dva/s) and durations of 40 ms (Collewijn, Erkelens, & Steinman, 1988). When trying to manipulate the color of the patch during the saccade, so that upon landing an updated stimulus with a new color is displayed to the observer, we as experimenters have to consider at least four additive sources of latency in our experimental setup (Figure 1) to be able to make the presentation deadline of each saccade offset.

**Figure 1.**
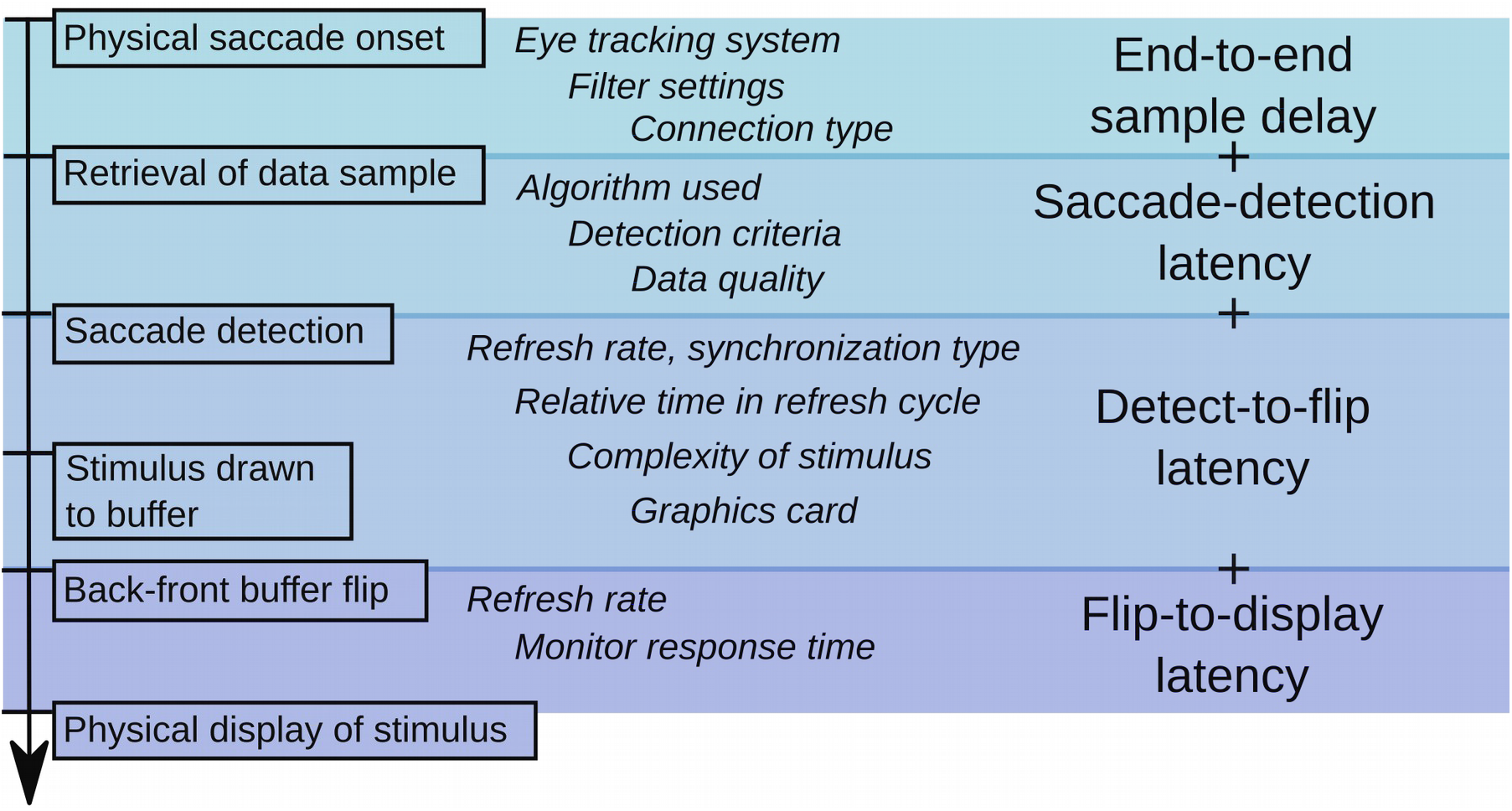
Schematic illustrating four categories of latencies in a temporal sequence (top to bottom) occurring when display changes are locked to saccade onset. Factors influencing the magnitude of the delays are shown in italics.

First, the online access to gaze position data is delayed. This *end-to-end sample delay* includes not only the time taken for a physical event to be registered, processed and made available online by the eye tracking system (e.g., capturing an image of the eye, fitting the pupil and corneal reflection, extrapolating gaze position), but also the time needed to retrieve that data via Ethernet, USB or analog ports. Although the retrieval time is usually negligible (i.e., on the order of μs), the total end-to-end sample delay can be considerable. According to manufacturer manuals, it may range from 1.8 - 3 ms in the Eyelink 1000 (SR-Research, 2010), from 1.7 - 1.95 ms in the Trackpixx3 (VPixx Technologies, 2017), from 3 - 14 ms in the Eyelink II (SR-Research, 2005), and up to 33 ms in the Tobii TX Series (Tobii Technology AB, 2010).

Second, as we need a reliable and thus often more conservative criterion to decide whether a saccade was actually initiated, the onset of the saccade detected online usually lags behind the offset of the saccade detected offline. Henceforth, this delay will be referred to as *saccade-detection latency*. Techniques to detect saccades during experiments often involve an invisible spatial boundary (Figure 2c) at some distance from the initial fixation point that gaze position has to cross (Rayner, 1975). This widely used technique (e.g., Collins, Rolfs, Deubel, & Cavanagh, 2009; Szinte & Cavanagh, 2011; Kalogeropoulou & Rolfs, 2017) usually provides reliable, but late saccade detection (~15 ms after actual saccade onset at a sampling rate of 500 Hz for a boundary 2 dva from fixation; Figure 5a). An alternative to the boundary technique is based on velocity thresholds (Figure 2d): The measured speed of the eye must exceed a certain value, such as 30 dva/s (Deubel, Schneider, & Bridgeman, 1996; Han, Saunders, Woods, & Luo, 2013), 40 dva/s (Castet, Jeanjean, & Masson, 2002) or even 100 dva/s (Arabadzhiyska, Tursun, Myszkowski, Seidel, & Didyk, 2017), detect saccades much earlier, but often suffer from increased false alarm rates.

**Figure 2.**
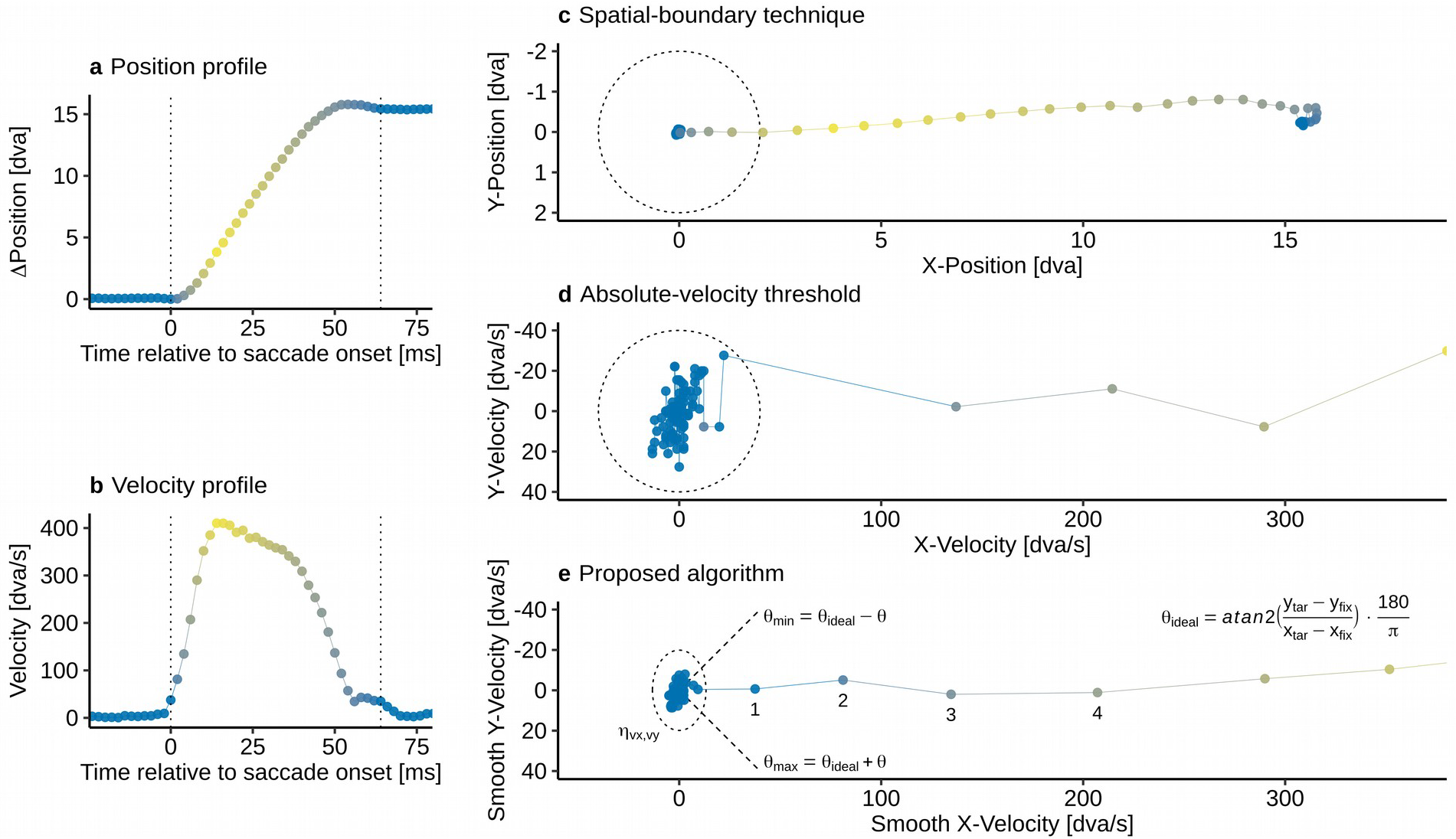
Illustration of different saccade detection techniques based on a exemplary saccade. **a-b** Plots of a position and velocity profile of a horizontal, rightward 15 dva saccade, sampled uniformly at 500 Hz. Color represents the sample-to-sample velocity (yellow: peak velocity). **c** Illustration of saccade detection using a spatial-boundary technique. Saccades are detected once gaze position reaches past the spatial boundary, defined by a 2 dva radius (dotted circle) around the instructed fixation position. **d** Illustration of an absolute-velocity threshold. Gaze position data is transformed into two-dimensional velocity space and a saccade is detected once velocities exceed a predefined value, e.g., 40 dva/s in this example. Once absolute velocity exceeds a defined value, here at 40 dva/s, a saccade is detected. **e** Illustration of the proposed algorithm. Gaze position data is resampled to a uniform sampling rate, transformed into two-dimensional velocity space, which is smoothed by a 5-point running average filter. Median-based standard deviations are computed separately for horizontal and vertical dimensions, forming an elliptic velocity threshold η_vx,vy_. An optional direction criterion θ (here 45°) can restrict detection to a range around the instructed saccade direction θ_ideal_ (e.g., computed via fixation and saccade landing positions xy_fix_ and xy_tar_) with θ_max_ and θ_min_ as upper and lower boundaries. The user may specify how many samples are needed that satisfy both velocity and direction criteria (here samples 1-4 are shown).

Third, once we have detected a saccade in the data retrieved online, the stimulus has to be drawn to the graphics card’s back-buffer and the flip with the front-buffer has to be synchronized with the display’s vertical retrace (Kleiner et al., 2007). This *detect-to-flip latency* is determined by the refresh rate of the monitor and depends on the time of detection within the refresh cycle. Novel technologies like G-Sync are able to reduce this latency to the sub-millisecond range by allowing flips as soon as rendering is complete without having to wait for the screen refresh (Poth et al., 2018).

Fourth, there is the *flip-to-display latency*, i.e., the time from the execution of the flip until the physical stimulus presentation on the screen. While the transfer of the entire video signal takes up to one frame duration, the display’s reaction time can additionally increase the flip-to-display latency, as well as introduce temporal jitter.

Taking into account all sources of delay (e.g., 5 ms end-to-end sample delay using an Eyelink II at 500 Hz with normal link filtering + 15 ms detection latency using a boundary technique + 5 ms mean detect-to-flip latency with a 120 Hz monitor + 8.3 ms flip-to-display latency), the physical change will occur in the last quarter of the 40 ms saccade. As both gaze-contingent displays and saccade profiles can be subject to considerable variance, we thus increase the risk of achieving a post-saccadic instead of the of intended intra-saccadic display change. Failure to acknowledge or control these latencies may thus lead to erroneous results and unwarranted conclusions.

While most latencies mentioned above largely depend on the specific hardware used, we can optimize the saccade-detection latency to achieve low-latency gaze-contingent presentations. Crucially, the choice of the saccade detection criterion determines both the timing and the reliability of the experimental paradigm: While a conservative detection criterion (e.g., a spatial boundary) may provide reliable, but late detection, a liberal detection criterion (e.g., a low absolute velocity threshold) may lead to early detection at the cost of increased false alarm rates. This may become especially relevant when detecting saccades based on velocity using high sampling frequencies, as any error in gaze position divided by a shorter sampling interval will lead to amplified velocity estimates (Han, Saunders, Woods, & Luo, 2013). To achieve reliable online detection, velocity thresholds would therefore have to be manually adjusted to precision and sampling frequency of the eye tracker, as well as to the situation- and participant-dependent noise levels (see also Engbert & Mergenthaler, 2006). To date, there is no algorithm that provides both fast and early online saccade detection while remaining reliable in noisier conditions at the same time.

Here we present a velocity-based online saccade detection algorithm that adaptively estimates noise levels based on preceding fixation data to provide robust results in the presence of random sample-to-sample noise, dropped samples, blinks, and fixational eye movements, while allowing its user to flexibly adjust the detection criterion to the specific experimental situation. We tested the performance of the algorithm and the impact of various parameter combinations and noise levels in a large-scale simulation with more than 34,000 saccades, and compared the algorithm to boundary techniques and absolute velocity thresholds. We then present a objective and perceptual test for reliable, gaze-contingent, and strictly intra-saccadic presentations that underlines the algorithm’s usefulness in real-time experimental scenarios.

### The algorithm

Online saccade detection relies on the continuous sampling of gaze position data (x, y) and the corresponding timestamps (t) throughout each trial of the experiment. Gaze position data collected during fixation is used to establish a threshold to demarcate the transition from fixation to saccade. Following Engbert & Kliegl’s widely used algorithm for microsaccade detection (Engbert & Kliegl, 2003; Engbert & Mergenthaler, 2006), the algorithm thus detects the onset of a saccade based on a elliptic, two-dimensional velocity threshold η_vx,vy_ (dotted line, Figure 2d), as defined by the product of the median-based standard deviation of horizontal and vertical gaze position dimensions (σ_vx_, σ_vy_) and a free scaling parameter λ to adjust the velocity criterion.

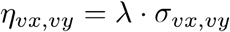

In addition, the user may provide a parameter *k* specifying how many of the most recent of all velocity samples must exceed the defined threshold. That way, robustness against false alarms due to noise-related velocity peaks is increased. In case the user intends to limit detection of saccades to an instructed saccade direction (θ_ideal_), which is often the case in controlled experimental paradigms, the algorithm allows for specification of an additional direction criterion θ that determines the direction range around the ideal saccade direction that individual velocity samples are allowed in (dashed lines, Figure 2d). This direction criterion can be used to avoid false detections of the instructed saccade as a consequence of other eye movements events that may satisfy the velocity criterion, such as blinks or microsaccades.

To make the algorithm suitable for online applications, two important features were implemented. First, owing to the fact that during online experiments it is rarely possible to retrieve every single data sample, missing position samples are linearly interpolated, either to a sampling rate specified by the user or to a sampling rate computed based on the number of samples retrieved in a given time. Second, two-point velocity samples are computed (to avoid edge velocities of zero) and then smoothed by a 5-point running average to reduce the impact of high frequency noise(Engbert & Mergenthaler, 2006). To not overestimate the first and most recent velocity samples, the vector edges are padded with repetitions of the first and the most recent samples, respectively. Subsequently, based on smoothed velocity samples, the median velocities (in most cases equaling zero) and the median-based standard deviations (σ_vx_, σ_vy_) are computed as described by Engbert and colleagues (Engbert & Kliegl, 2003; Engbert & Mergenthaler, 2006; Engbert, Rothkegel, Backhaus, & Trukenbrod, 2016):

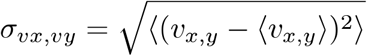

Brackets 〈.〉 stand for the median estimator. To optimize processing speed, we use the quick select algorithm for median selection (Press, Teukolsky, Vetterling, & Flannery, 2007).

To determine whether a saccade is ongoing, only the most recently retrieved *k* samples (*k* has to be defined by the user beforehand) are tested whether eye velocity exceeds the specified threshold, which is computed based on all preceding *n-k* samples. An ongoing saccade is detected only if all *k* samples pass this velocity test criterion *vel*:

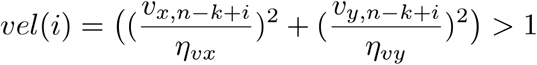

As mentioned above, in case of the application of an additional direction criterion *dir*, the direction of the same samples must also fall within a direction range specified by the user.

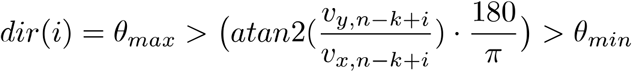

Based on this equation, the ideal saccade direction can be conveniently computed using the instructed fixation and saccade target regions (θ_ideal_, Figure 2d).

The algorithm automatically returns the used velocity thresholds, as well as (optionally) interpolated position data, and—if a saccade has been detected—the timestamp and computed eye velocity at detection. As online saccade detection by definition occurs after saccade onset and lower detection threshold are more susceptible to noise, the algorithm also provides an estimate for the actual saccade onset by tracing back in time one sample that falls below another velocity threshold, i.e., the product of user-defined threshold parameter λ_onset_(not necessarily the same as λ used for saccade detection) and the computed median-based standard deviation σ_vx,vy_. This two-step procedure (Dorr, Martinetz, Gegenfurtner, & Barth, 2010; Arabadzhiyska, Tursun, Myszkowski, Seidel, & Didyk, 2017) allows the user to get real-time access to a reliable timestamp of saccade onset, for instance to provide feedback on saccade latency in a certain trial, to trigger a display change at a pre-defined time relative to saccade onset, or to fit ongoing saccade trajectories (Han, Saunders, Woods, & Luo, 2013).

To code is openly available online at *https://github.com/richardschweitzer/OnlineSaccadeDetection*. It uses standard C libraries and can thus be used across platforms. We provide a module in Python, as well as an implementation to be compiled as a mex-function in Matlab (Mathworks, Natick, MA, USA).

## Materials and methods

### Simulation

#### Data

For the validation of the algorithm, we compiled a data set consisting of a total of 34607 saccades, measured from participants’ dominant eye. The data was collected from two past experiments (i.e., Schweitzer & Rolfs, 2017; Watson, Schweitzer, Castet, Ohl, & Rolfs, 2017), as well as from one pilot study. Using an Eyelink II at a sampling rate of 500 Hz, a number of 17501 horizontal (left- and rightward) saccades with an instructed amplitude of 14.6 dva, as well as a number of 10809 saccades in eight different directions (cardinal and intercardinal directions) and of 10 dva amplitude, entered analysis. Furthermore, collected with an Eyelink 1000+ at a sampling rate of 1000 Hz, we included 6297 additional saccades in the same eight directions, but of 8 dva amplitude.

Pre-processing of the data used for the validation (offline data analysis) involved three steps. First, trial data was reduced to those samples between the onset of successful fixation (preceding the saccade go signal) and 100 ms after the participant’s gaze first reached the target area (boundary with 2 dva radius around saccade target). Second, the onset and offset of the saccade - defined as the ground truth in all analyses - was detected using the Engbert-Kliegl algorithm (Engbert & Kliegl, 2003; Engbert & Mergenthaler, 2006) with a threshold parameter of λ = 5 and a minimum duration of 16 milliseconds (8 samples at 500 Hz and 16 samples at 1000 Hz). Trials, in which saccades could not be detected or in which more than one saccade occurred within the chosen time interval were excluded. Third, eye movement data was transformed from the setup-specific pixel values to degrees of visual angle. Subsequently, position data and timestamps were normalized relative to the detected onset of the saccade to allow for comparisons between saccades of different amplitudes and durations. Saccade data and code used for simulations are available on the Open Science Framework: https://osf.io/3pck5/.

#### Procedure

To simulate the performance of the online detection algorithm, we divided the data of each trial in saccade absent (i.e., prior to saccade onset as detected offline) and saccade present (i.e., after saccade onset as detected offline) segments. The algorithm was then run sequentially on each data sample (ordered by time stamps) in the respective segment, taking into account all previous samples for threshold estimation. That way, we simulated its usage during an experimental trial in which new data samples are retrieved cumulatively. If saccades were detected while iterating through absent segments, we registered a false alarm (FA), if not, the trial counted as a correct rejection (CR). Similarly, if saccades were detected after offline-detected saccade onset, we registered a correct detection (hit), if not, the trial counted as a miss. To evaluate the performance of the boundary technique (2 dva) and absolute velocity threshold (40 dva/s), we used the same procedure.

To explore the behavior of the algorithm in a larger parameter space, online saccade detection was tested in both absent and present segments for every available parameter combination, i.e., threshold parameter λ (levels: 5, 10, 15, 20), samples above threshold needed *k* (levels: 1, 2, 3, 4), and direction criterion θ (levels: none, 45°, 30°, 15°). In addition, we convoluted all data samples with Gaussian noise (SDs: 0, 0.025, 0.05, 0.1 dva) on both horizontal and vertical dimension and randomly removed a proportion of all samples (levels: 0, 10, 20, 30%), to simulate eye tracker noise and sample loss, respectively. This test setup resulted in a total of 1024 within-saccade conditions.

For each within-saccade condition and additionally for each available sampling rate and saccade direction, we computed detection sensitivity index *d*’ and summary statistics for detection latency (saccade present segments only) of the three detection methods, i.e., boundary techniques, absolute velocity thresholds, and the described online detection algorithm. In addition, we computed an efficiency score (ES), i.e., the proportion of correct rejections divided by the mean detection latency relative to the actual saccade onset (Townsend & Ashby, 1983).

#### Analysis

As a first step, we computed summary statistics (mean, standard deviation, standard error) for detection latency (separately for each online detection technique and within-saccade condition), and median-based standard deviation of velocity samples. For detection accuracy, we computed *d*’, proportion of hits and false alarms for each online detection technique and condition, and estimated their standard error using non-parametric bootstrapping with 2000 repetitions.

To understand the individual effects of the algorithm’s parameters on the dependent variables *d*’ and detection latency (ms), we applied multiple regression on the aggregated data. Sampling rate was included as an effect-coded factor (−0.5: 500Hz, +0.5: 1000 Hz), while threshold factor λ and samples above threshold *k* were included as continuous predictors centered around their mean. Direction restriction was also included as a continuous predictor (in degrees: 180, 45, 30, 15).

To analyze the online detection algorithms’ robustness against noise, we ran a second multiple regression on detection accuracy (d’) and detection latency (ms), including four factors and their interactions: sampling rate (effect-coded; −0.5: 500 Hz, +0.5: 1000 Hz), Gaussian noise standard deviation (continuous; 0, 1.5, 3, 6 arcmin), percentage of samples dropped (continuous; 0, 10, 20, 30%), and detection technique used (dummy-coded; boundary, absolute velocity, algorithm [λ=5], algorithm [λ=10], algorithm [λ=15], algorithm [λ=20]).

### Experimental Test

#### Participants

Ten observers (including the first author) participated in the experiment. All observers (four female; age range: 22 - 35 years old) had normal or corrected-to-normal vision. The study was conducted in agreement with the Declaration of Helsinki (2008), approved by the Ethics Committee of the German Society for Psychology, and all observers provided written informed consent before participation. We tracked participants’ dominant eye (eight of ten observers with right ocular dominance) for one session with an average duration of 30 minutes for 480 trials in total.

#### Apparatus

The experiment took place in a dimly lit, sound-attenuated cabin. A Propixx DLP Projector (Vpixx Technologies, Saint-Bruno, QC, Canada) running at a temporal resolution of 1440 frames per second and a spatial resolution of 960 x 540 pixels projected into the cabin onto a 200 x 113 cm screen (Celexon HomeCinema, Tharston, Norwich, UK). The projector was connected to the experimental host-PC via a Datapixx3 (Vpixx Technologies, Saint-Bruno, QC, Canada). Observers were seated at a distance of 180 cm away from the projection screen with their head supported by a chin rest. Stimulus display was controlled using the PsychProPixx function from PsychToolbox (Pelli, 1997; Kleiner et al., 2007) running in Matlab 2016b (Mathworks, Natick, MA, USA) on a custom-build desktop computer with an Intel i7-2700K eight-core processor, 8 GB working memory, and a Nvidia GTX 1070 Ti graphics card, running Ubuntu 18.04.1 (64-bit) as operating system. The setup is illustrated in Figure 3. Eye tracking was performed using an Eyelink 1000+ desktop base system, tracking participants’ dominant eye at a sampling rate of 2000 Hz. Tracking was controlled during the experiment using the Eyelink Toolbox (Cornelisen, Peters, & Palmer, 2002). Moreover, we collected data from a photodiode connected to an actiChamp EEG amplifier (Brain Products, Gilching, Germany), that was attached to the lower right corner of the projection screen (i.e, at approximately 36 dva eccentricity relative to central fixation), again at a sampling rate of 2000 Hz. To synchronize eye tracking and photo sensor data, we applied a DB-25 Y-splitter cable to simultaneously send triggers of 1 ms duration to the Eyelink host computer and EEG host computer. During pre-processing of the data, we used the EYE-EEG Toolbox (Dimigen, Sommer, Hohlfeld, Jacobs, & Kliegl, 2011) in EEGLAB (Delorme & Makeig, 2004) to temporally align both recordings. For all triggers across recordings, we found a mean absolute misalignment error of 0.38 ms, i.e., below one sample.

**Figure 3.**
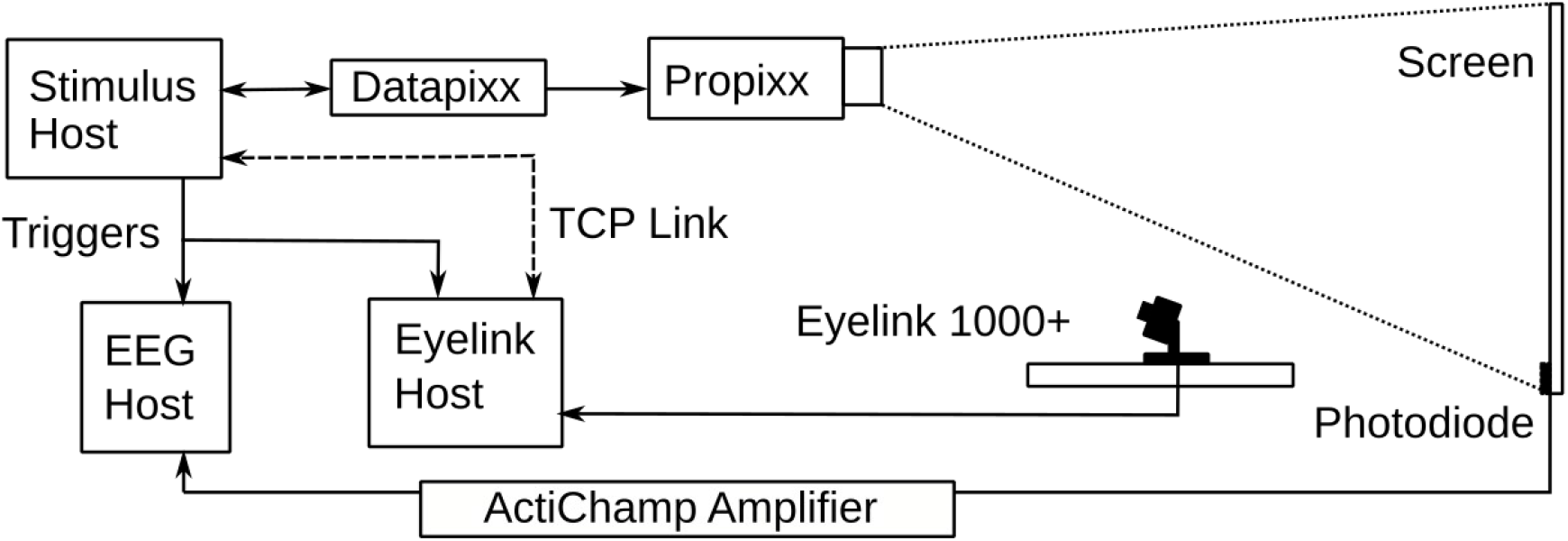
Setup used to co-register gaze position and photodiode data. The stimulus host computer performed online monitoring of gaze position via the TCP link, stimulus presentation using a ProPixx DLP projector with a frame rate of 1440 Hz, as well as synchronized triggering of EEG and Eyelink host computers, recording photodiode and gaze position, respectively.

#### Stimuli

Stimuli were Gabor patches of vertical orientation enveloped in a Gaussian window with a standard deviation of 0.5 dva, presented on a uniform grey background (luminance of 30 cd/m^2^). All Gabor patches had a spatial frequency of 0.5 cycles per degree of visual angle (cpd) and a contrast of 100% (0% in stimulus absent conditions).

In both saccade and fixation trials (see below and Figure 4), stimulus presentation duration amounted to 20 frames at a frame rate of 1440 Hz, that is, 13.9 ms. In order to reduce the transient elicited by a sudden stimulus onset, the first four and last four presentation frames were used to linearly ramp up and down stimulus contrast, respectively. Presentation locations were randomly chosen in each trial: Relative to the screen center, stimuli could appear at an eccentricity of up to 8 dva within a range of 360 degrees. As we aimed to present stimuli at largely the same retinal eccentricities both during saccade and fixation trials, we estimated that intra-saccadic presentations would be realized in the first half of the saccade and would therefore be effective when the saccade crossed the screen’s center vertical midline. In fact, across all participants’ gaze position at stimulus onset was 1.24 dva (SD = 0.92 dva) left of the vertical midline for rightward saccades and 1.31 dva (SD = 1.0 dva) right of it for leftward saccades.

**Figure 4.**
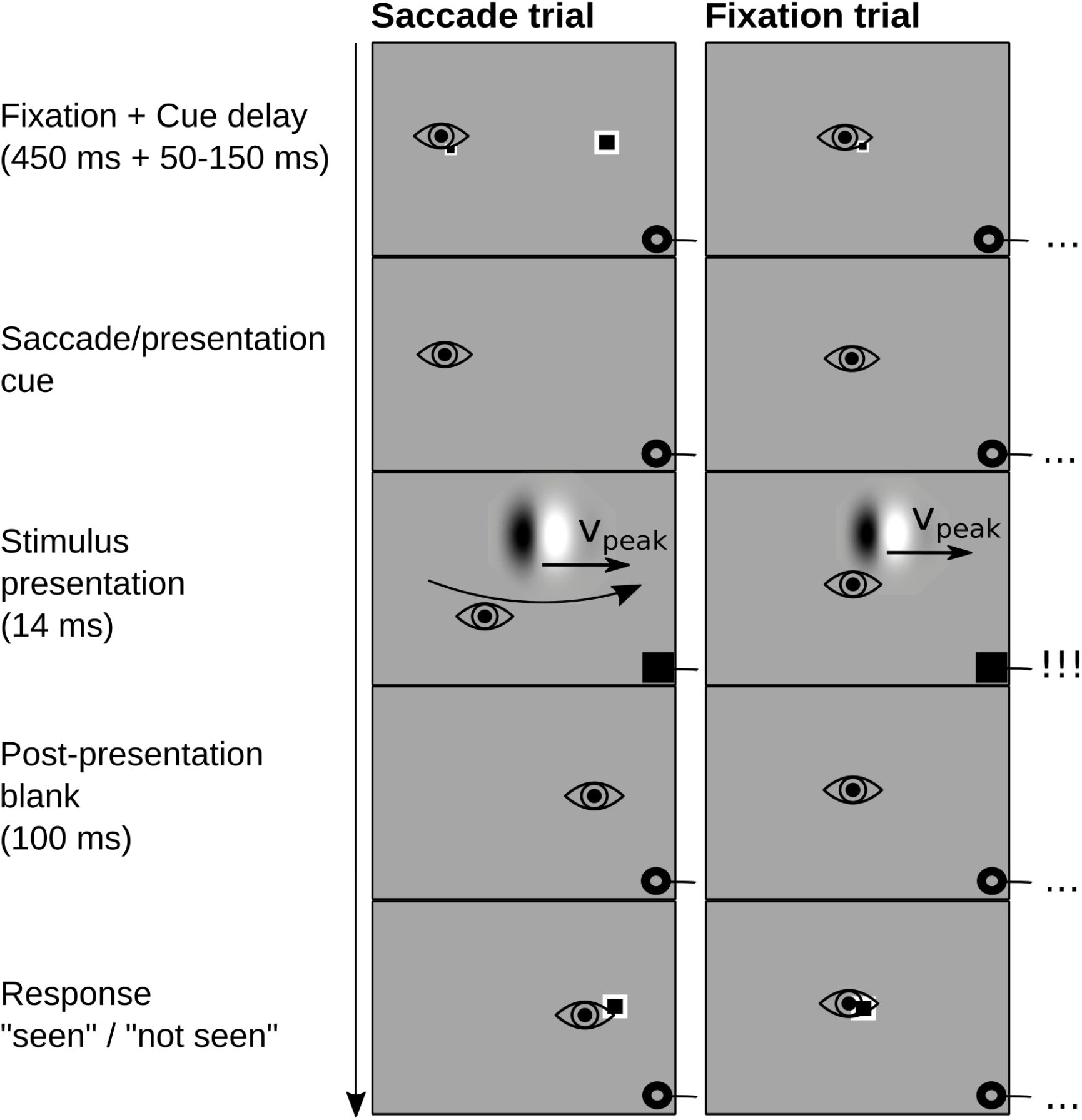
Experimental procedure used in saccade and fixation trials. In saccade trials, observers made a 16 dva saccade, whereas in fixation trials, observers maintained central fixation. In saccade trials as soon as a saccade was detected online and in fixation trials after the observer’s median saccade latency, a Gabor patch (vertical orientation, 0.5 cpd) enveloped in a Gaussian window with a standard deviation of 0.5 dva and drifting at saccadic peak velocities (median peak velocity of 30 most recent saccade trials) was presented for 13.9 ms. During stimulus presentation, a black square was projected onto a photodiode located in the lower right corner of the screen, generating a signal change in the photodiode.

Throughout presentation, Gabor patches were drifting at constant speeds equivalent to saccadic peak velocities, which were automatically computed during the experiment. That is, after each saccade trial, gaze position data collected during the trial was resampled to 500 Hz and cropped to the relevant time interval between cue onset and 30 milliseconds after reaching the target region. Second, we used the Engbert-Kliegl saccade detection algorithm (Engbert & Kliegl, 2003; Engbert & Mergenthaler, 2006) with a minimum duration of 8 samples and λ = 10 to extract saccade latency and saccadic peak velocity. Third, we computed the median saccadic peak velocity based on the 30 most recent saccade trials. Fourth, to investigate the effect of stimulus drift velocity relative to saccade velocity, we defined the stimulus drift velocity as the resulting median or added or subtracted 50 dva/s, based on which we then computed the Gabor’s phase change per frame. This procedure resulted in three conditions and distributions of stimulus drift velocities (M_−50_ = 366 dva/s, M_0_=416 dva/s, M_+50_=466 dva/s), matching the mean saccadic peak velocity of 419 dva/s. Although in both conditions drift velocity was computed based on the most recent saccadic peak velocities, given the fact that fixation and saccade trials were presented in an interleaved manner, presented drift velocities might have differed between conditions. This, however, was not the case (t_−50_(9)=−0.28, p_−50_=.80; t_0_(9)=−1.8, p_0_=10; t_+50_(9)=−0.43, p_+50_=.68). To achieve visibility during saccades, Gabor patches always drifted in the direction of the saccade (Castet & Masson, 2000; Deubel, Elsner, & Hauske, 1987).

The instructed fixation location was marked using a full-contrast black rectangular dot with a white outline and size of 0.4 dva. For the saccade target location, a similar stimulus was applied, only of twice the size, i.e., 0.8 dva.

#### Procedure

Each participant ran a total of 480 trials, consisting of 240 saccade trials and 240 fixation trials. For each trial type, there were 120 stimulus-absent and 120 stimulus-present trials, which then contained three stimulus velocity conditions (sum of median peak velocity and either −50, 0, or +50 dva/s) and two stimulus drift directions (leftward vs. rightward, in saccade trials according to saccade direction). All trials were presented in interleaved and randomized order.

##### Saccade trials

Each saccade trial (Figure 4, left column) began with the display of two dots (see *Stimuli*), of which the smaller one represented the fixation location and the larger one the saccade target. Both dots had a horizontal eccentricity of 8 dva relative to the screen center. After successful fixation within a 2 dva radius around the fixation dot for 450 ms, followed by a random delay of 50 to 150 ms, both dots disappeared from the screen, i.e., the saccade cue. Participants were instructed to make a saccade (16 dva) towards the remembered target location right after the disappearance of the dots. Saccades were detected online within a window of 10 s (mean saccade latency was 275 ms, SD = 135 ms) after the onset of the saccade cue using the algorithm described in this paper (parameters: λ = 10, k = 3, θ = 30°). As soon as a saccade in the instructed direction was detected, we triggered the presentation of a Gabor patch drifting at saccadic peak velocities. In stimulus-present trials, the patch occurred intra-saccadicly for 13.9 ms within a radius of 8 dva around the screen center with 100% contrast, whereas in stimulus-absent trials, the patch had zero contrast. Stimulus-absent and -present trials were present in an interleaved manner and were equally probable. Regardless of whether a stimulus was present or absent in a given trial, the presentation was always accompanied by a black dot with a size of 4 dva which was displayed (for the same time as the stimulus) at the location of the photodiode attached to the lower right corner of the screen. 100 ms after stimulus offset (i.e., on average 82 ms after saccade offset), the saccade target dot would reappear to give participants feedback on the accuracy of their saccade and prompt their response. Participants were instructed to respond with RightArrow in case they detected a stimulus and LeftArrow in case they did not. They did not receive feedback on their detection performance, but were shown their own saccade trajectory on the screen whenever they did not reach the saccade-target area (2 dva radius) or made more than one saccade before reaching the latter. Trials with these insufficient saccades were not repeated.

##### Fixation trials

Fixation trials (Figure 4, right column) were initiated with the display of a small dot (0.4 dva) representing the center of a fixation area with 2 dva radius. Just like during saccade trials, gaze had to stay within this area for 450 ms to initiate the presentation sequence (plus random delay of 50 to 150 ms), until the dot disappeared. Prior to stimulus presentation, a delay with the duration of the participant’s median saccade latency (based on the 30 most recent saccade trials) was added to imitate the saccade trials and to increase temporal predictability. For the presentation of the rapidly drifting Gabor patch and the photodiode dot under fixation, the same parameters were applied as during saccade trials (see *Stimuli* and *Saccade trials* above). Again, 100 ms after stimulus offset a larger dot (0.8 dva) appeared, prompting the participant’s response (RightArrow: “seen”, LeftArrow: “not seen”). Participants were instructed to maintain fixation until the appearance of the response cue and received feedback whenever their gaze left the fixation area.

#### Analysis

We collected a total of 480 trials (240 saccade trials and 240 fixation trials in interleaved and randomized order) per participant plus one additional set of 305 trials from one participant owing to an aborted session. Due to insufficient fixation or early responses, 9.2% of all fixation trials had to be excluded. In saccade trials, 17.1% were excluded due to unsuccessful initial fixations, not reaching the target area with only one saccade or responding before having reached the target area. Although the Gabor stimuli should be invisible during fixation due to their high drift velocity (Castet & Masson, 2000; Deubel, Elsner, & Hauske, 1987; Garcìa-Pérez & Peli, 2011; Watson, Schweitzer, Castet, Ohl, & Rolfs, 2017), one percent of the remaining saccade trials were still excluded because the saccade offset preceded the stimulus offset (as measured by the photodiode). Finally, 0.4% of all trials were removed because of dropped frames. On average, 222 (SD = 13) of the initial 240 fixation trials and 201 (SD = 23) of the initial 240 saccade trials were included for analysis.

Photodiode data and eye movement data were merged using the EYE-EEG Toolbox (Dimigen, Sommer, Hohlfeld, Jacobs, & Kliegl, 2011) and downsampled to 1000 Hz. Saccades were detected using Engbert-Kliegl algorithm (Engbert & Kliegl, 2003; Engbert & Mergenthaler, 2006) with a threshold of λ = 5 and a minimum saccade duration of 16 samples, constituting the ground truth for the analyses on latency. In addition, messages on saccades and fixations generated by the Eyelink system were imported to validate the saccade detection results from the Engbert-Kliegl algorithm. Unfiltered photodiode voltage time series data was transformed to standard z-scores separately for each experimental session, so that the standard luminance of the screen produced values around 0 and the reduction in photodiode response due to the brief presentation of the black dot during stimulus presentation resulted in values well below −4. To determine whether the stimulus was on the screen we thus selected those values below the cutoff of −3. In both saccade and fixation trials, we computed retinal velocity of the stimulus during its presentation by estimating eye velocity (using a five-point running mean) from those gaze samples collected during stimulus presence as determined by the photodiode, and subtracting it from the drift velocity of the stimulus.

To analyze the influence of retinal velocity on task performance on a trial-by-trial basis, we used a logistic mixed-effects regression with random intercepts and slopes for observers (Bates, Mähler, Bolker, & Walker, 2015) to predict correct responses from retinal velocity (continuous predictor, z-scores computed separately for fixation and saccade conditions) and condition (effect-coded; −0.5: Fixation, +0.5: Saccade). Confidence intervals for log odds weights were determined via parametric bootstrapping with 10,000 repetitions. Hierarchical model comparisons were performed using the likelihood ratio test.

We furthermore used t-tests to determine whether task performance (d’) was different from chance levels and a univariate repeated-measures ANOVA to compare task performance in fixation and saccade conditions.

## Results

### Simulation results

Here we simulated a situation similar to most experimental paradigms: Once an observer receives a cue to make a saccade, we continuously retrieve data samples from the eye tracker to determine whether at any point in time the observer has initiated a saccade or not. In this setup, detection performance has two main aspects, namely accuracy (i.e., detection after saccade onset and not before that) and latency (i.e., when is a saccade detected relative to the actual saccade onset, as determined offline). In this simulation we asked two main questions: (1) How is the performance of the proposed algorithm impacted by the choice of parameters and (2) how does its performance compare to classic techniques, such as spatial boundaries and absolute velocity thresholds, especially under conditions of additional noise and data loss?

As shown in Figure 5a, online saccade detection is a trade-off between speed and accuracy. At a sampling rate of 500 Hz, for instance, boundary techniques (red squares), have a very high accuracy (p(FA) = 0.4%, SD = 0.3%; mean d’ = 6.3, SD = 0.74), but long saccade-detection latencies (M = 15 ms, SD = 1.1 ms), whereas absolute velocity thresholds (green squares) have shorter detection latencies (M = 4.4 ms, SD = 0.23 ms), however with lower accuracy (p(FA) = 11.6%, SD = 3%; mean d’ = 4.6, SD = 0.34). Importantly, the type of eye tracking system and the sampling frequency of the eye tracker are major moderators of the performance of both techniques. At low sampling rates, samples become less frequently available, whereas at high sampling rates sample-to-sample noise impacts velocity estimates to a larger extend (Han, Saunders, Woods, & Luo, 2013). For comparison, at a sampling rate of 1000 Hz, saccade-detection latencies of both techniques decrease as compared to 500 Hz (boundary: M = 11.2 ms, SD = 0.96 ms; absolute velocity: M = 1.4 ms, SD = 0.08 ms), whereas false alarm rates increase drastically for absolute velocity thresholds (p(FA) = 63%, SD = 6%; mean d’ = 2.8, SD = 0.16). In contrast, detection accuracy of the proposed algorithm (shapes in shades of violet) remained largely unaltered across sampling frequencies (500 Hz: mean p(FA) = 11.7%, SD = 2.5%; d’ = 5.1, SD = 1.0; 1000 Hz: mean p(FA) = 14.6%, SD = 3.2%; d’ = 5.1, SD = 1.5; β = −0.01, t(128) = −0.06, p = .95), due to the adaptive adjustment of the noise level, while detection latency decreased (500 Hz: M = 5.5 ms, SD = 2.1 ms; 1000 Hz: M = 3.7 ms, SD = 1.9 ms; β = −1.5, t(128) = −6.1, p < .0001).

**Figure 5.**
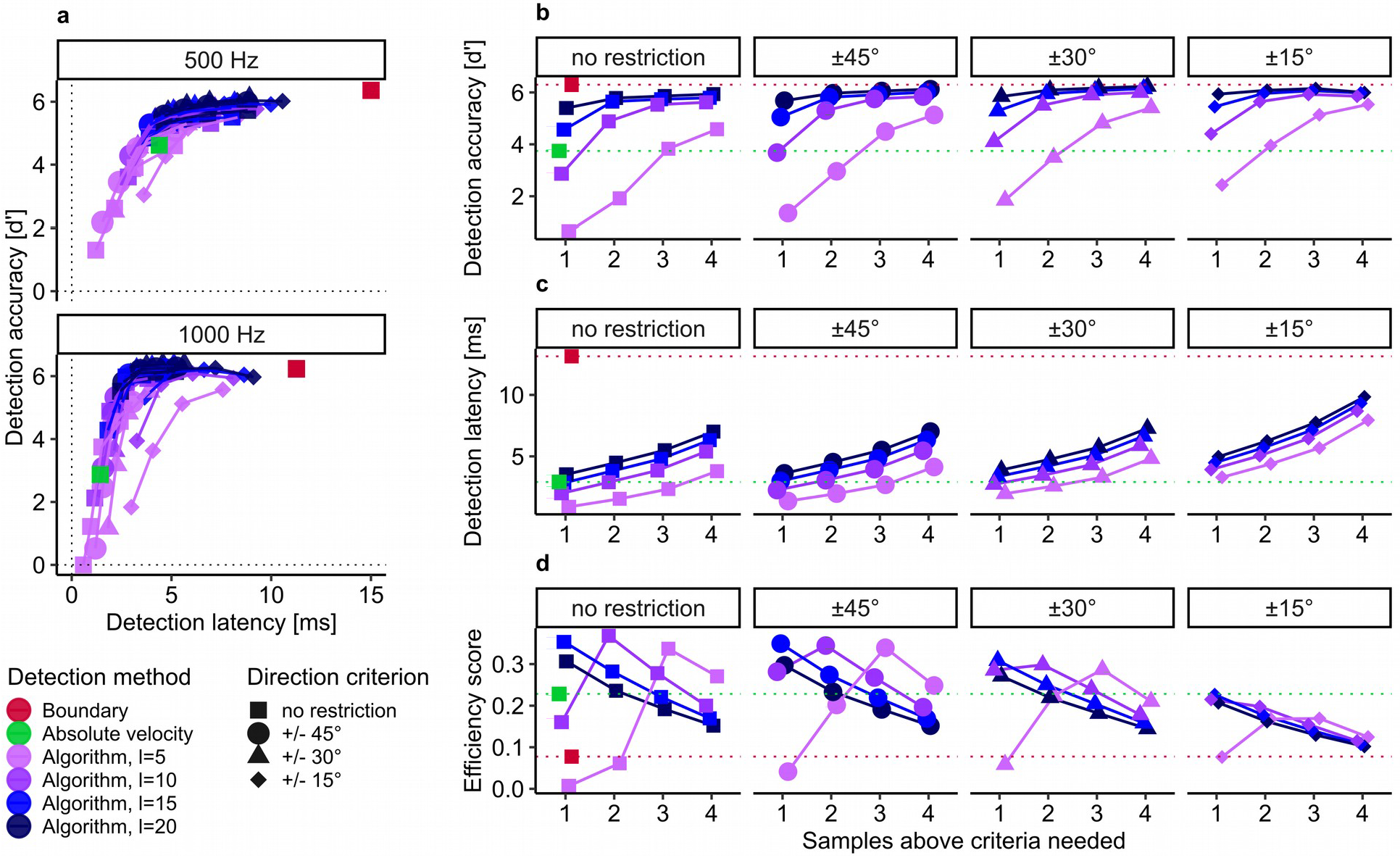
Grand averages of detection performance and latency, as determined by simulation. **a** The trade-off of detection accuracy and detection latency for each sampling rate. Every dot represents the mean across all trials including all eight tested saccade directions, color indicating the type of detection method (and threshold factor λ) used, shape indicating the direction criterion (θ) used. The four connected values indicate the number of samples above threshold (k) needed for detection in each condition (always increasing from left to right). **b-d** Mean detection accuracy, latency, and efficiency averaged across sampling rates for different parameter combinations (λ, θ, k). Green and red dotted reference lines indicate the average values for absolute velocity thresholds and boundary techniques, respectively.

Saccade-detection accuracy (Figure 5b) and latency (Figure 5c), however, strongly depend on the choice of the necessary parameters *k*, λ, and θ. First, increasing the number of samples needed above threshold *k* improved accuracy (d’) by 0.36 per sample (β = 0.36, t(128) = 4.3, p < .0001), but also increased latency by 1.38 ms per sample (β = 1.38, t(128) = 8.9, p < .0001). Second, a higher threshold parameter λ similarly increased both accuracy (β = 0.1, t(128) = 6.1, p < .0001) and latency (β = 0.18, t(128) = 5.8, p < .0001). Third, accepting a wider range of saccade directions (in degrees) led to a decrease of accuracy (β = −0.004, t(128) = −5.9, p < .0001) and latency (β = 0.001, t(128) = −6.9, p < .0001). While for saccade-detection latency all three parameters had additive effects (Figure 5c), there were interactions present for detection accuracy: The benefit of increasing the number of samples above criteria *k* (β = −0.07, t(128) = −6.6, p < .0001) or restricting directions *θ* was smaller at higher thresholds (β = 0.0004, t(128) = 3.2, p = .002), because detection accuracy would reach ceiling (Figure 5b). Accordingly, direction restriction was more effective at low λ and low *k* (β = 0.001, t(128) = 2.1, p = .043).

To improve the understanding of this speed-accuracy tradeoff, we introduced an efficiency score (Townsend & Ashby, 1978) based on the ratio of correct rejection rate and detection latency (Figure 5d). It becomes evident that for optimal parameter choice, the efficiency of the proposed algorithm is well above the efficiency of both boundary and absolute velocity techniques. With extremely conservative settings (see Figure 5b-d, rightmost panel), however, detection latency will be increased to a large degree, so that some saccades might not be detected in time. With regard to the optimal choice of parameters, it is important to consider noise levels and sampling rate of the eye tracker. For our simulation, we chose two eye trackers with similar spatial precision (RMS = 0.01 dva; SR-Research, 2005; SR-Research, 2010), but with varying sampling rate. We found that the positive effect of detection threshold λ and number of samples above threshold *k* on detection accuracy was slightly stronger when tracking at higher than at lower sampling rates (λ: β = 0.047, t(128) = 1.99, p = .049; *k*: β = 0.24, t(128) = 2.01, p = .038). This suggests that a more conservative parameter choice is more beneficial at higher sampling rates, where increased velocity due to tracker noise is more likely to occur (see also Figure 6, bottom row).

**Figure 6.**
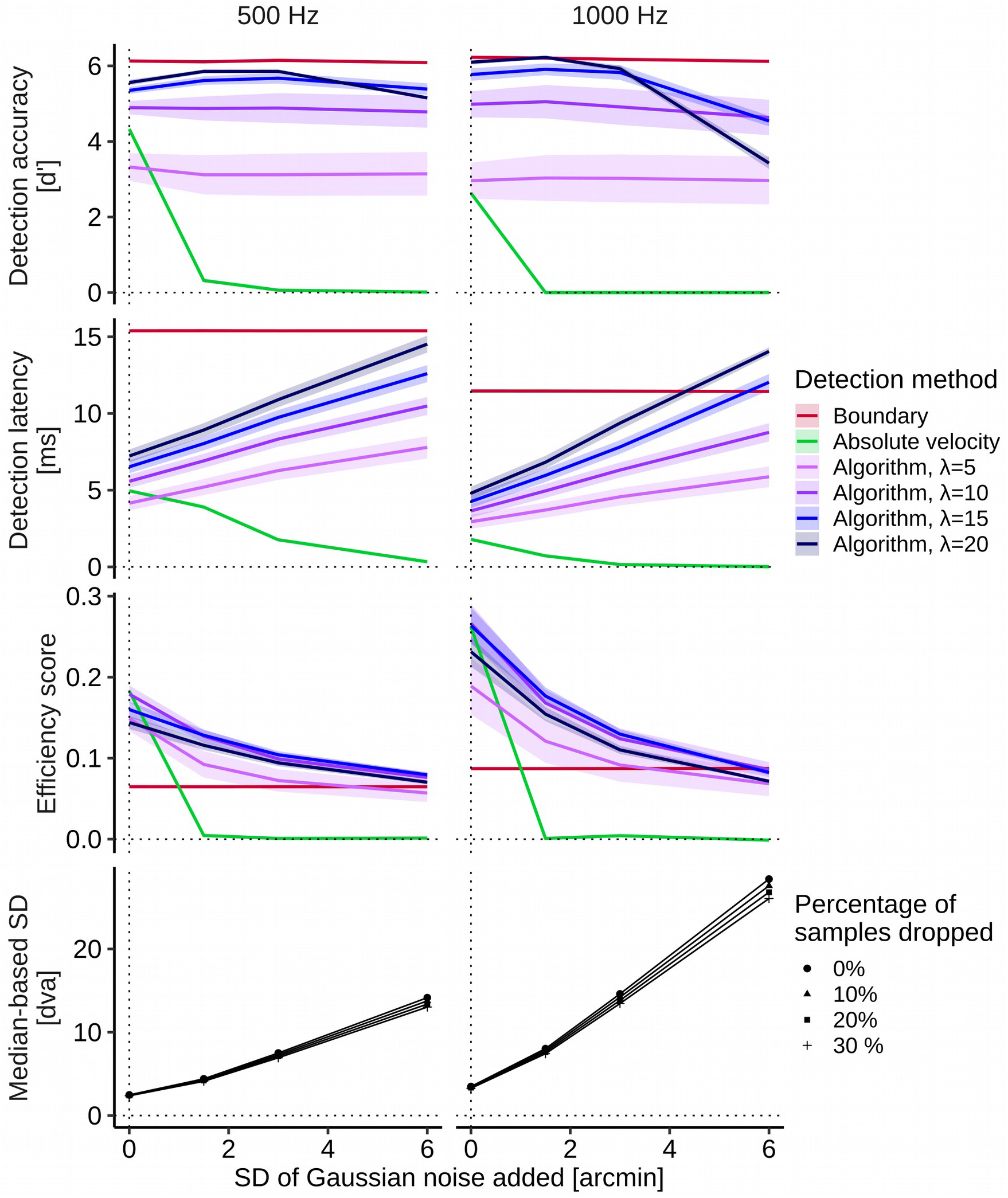
Mean detection accuracy, latency, and efficiency of the three online saccade detection techniques for different noise levels (SD of Gaussian noise added to both sample dimensions, x and y) and sampling rates (left column: 500 Hz, right column: 1000 Hz) averaged across all levels of percentage of samples dropped. Shaded areas indicate 95% confidence intervals. **Bottom row:** Median-based standard deviations of absolute velocity estimates used to compute velocity thresholds.

How do online saccade detection techniques cope with conditions in which noise is drastically increased or in which several samples are dropped? We simulated these situations by adding uncorrelated, Gaussian noise (standard deviations of up to 6 arcmin) to the data and by randomly removing data samples (up to 30%). As shown in Figure 6 (top row), absolute velocity thresholds (green lines) are strongly impacted by noise (β = −0.49, t(144) = −4.7, p < .0001), as the false alarm rate reached almost 100 % after adding Gaussian noise of 1.5 arcmin SD, effectively reducing this technique’s efficiency to virtually zero, i.e., 0.005 (SD = 0.001) at 500 Hz and 0.001 (SD = 0.0002) at 1000 Hz. Detection accuracy of the proposed algorithm (500 Hz: mean efficiency = 0.11, SD = 0.036; 1000 Hz: mean efficiency = 0.15, SD = 0.016), on the other hand, was largely unimpaired by noise (λ=5: β = −0.02, t(144) = −0.15, p = .88; λ=10: β = −0.04, t(144) = −0.36, p = .72; λ=15: β = −0.11, t(144) = −1.10, p = .27). At a threshold factor of λ=20, however, detection accuracy decreased starting at Gaussian noise SDs of 6 arcmin (β = −0.29, t(144) = −2.75, p = .007), since velocity thresholds were simply too high: If median-based velocity SDs such as 26 dva/s (Figure 6, bottom row, right panel) were multiplied by a factor of 20, we would achieve unreasonable velocity thresholds as high as 520 dva/s and thus miss most ongoing saccades.

As the velocity threshold is estimated based on the current noise level to preserve robustness across trials and participants, higher velocity thresholds due to increased noise levels should be accompanied by increased detection latencies. Indeed, for every threshold factor λ, latency increased with noise level (λ=5: β = 0.55, t(144) = 14.3, p < .0001; λ=10: β = 0.85, t(144) = 22.12, p < .0001; λ=15: β = 1.19, t(144) = 30.9, p < .0001; λ=20: β = 1.42, t(144) = 36.7, p < .0001). While being sensitive to changes in the noise level, the algorithm was robust against dropped samples, with respect to both accuracy (λ=5: β = −0.01, t(144) = −0.63, p = .53; λ=10: β = −0.005, t(144) = −0.28, p = .77; λ=15: β = −0.002, t(144) = −0.1, p = .92; λ=20: β = −0.001, t(144) = −0.01, p = .99) and latency (λ=5: β = −0.003, t(144) = −0.43, p = .66; λ=10: β = −0.001, t(144) = −0.1, p = .92; λ=15: β = −0.0003, t(144) = −0.05, p = .95; λ=20: β = −0.002, t(144) = −0.28, p = .78). As a comparison, both boundary techniques and absolute velocity thresholds suffered from longer detection latencies due to dropped samples (boundary: β = 0.021, t(144) = 4.2, p = .0004; absolute velocity: β = 0.014, t(144) = 1.98, p = .049).

In a separate simulation, we found that the algorithm’s running time (i.e., the time elapsed from invocation to return of the mex-function in Matlab 2016b on a Dell Optiplex 3020 with an Intel i5-4590 processor running Kubuntu 18.04) on data collected for two seconds was on average 0.051 ms at sampling rate of 500 Hz (SD = 0.005 ms, N = 30000), 0.097 ms at 1000 Hz (SD = 0.009 ms, N = 30000), and 0.187 ms at 2000 Hz (SD = 0.011 ms, N = 30000). The algorithm’s time average complexity is thus linear (and quadratic in the worst case).

### Experimental results

To present an example for the application of the algorithm and to show that its application make strictly intra-saccadic presentations well possible, we developed an objective and a perceptual test. As our setup allowed for the co-registration of stimulus events, EEG recordings and eye tracking at a high temporal resolution, the objective test used a photodiode to measure physical stimulus onset and offset during the saccade, where the saccade was detected online with the proposed algorithm (see Figure 3 for Apparatus). In addition, as a perceptual test, we presented a Gabor patch (vertical orientation, 0.5 cpd spatial frequency, 0.5 dva SD Gaussian aperture) that – due to its high drift velocity (on average 420 dva/s) – would be invisible during fixation, but become detectable only if the eye was moving at a similar velocity in the same direction (Castet & Masson, 2000; Watson, Schweitzer, Castet, Ohl, & Rolfs, 2017). Observers were instructed to indicate whether they perceived a Gabor patch that could be presented anywhere within a range of 8 dva around screen center and was present in 50% of all saccade and fixation trials (Figure 4).

Online saccade detection in saccade trials worked well. Only 1% of valid trials had to be excluded due to too early or too late detection. Mean detection latency amounted to 5 ms (SD = 2.06 ms). Unlike results from the simulation, however, this latency estimate still included system delays, most crucially the end-to-end sample delay of the eye tracker: According to the manufacturer, the Eyelink 1000+ with normal filter settings at a sampling rate of 2000 Hz is expected to have an average delay of 2.7 ms (SD = 0.2 ms; SR-Research, 2013). Correcting for these delays, the mean detection latency would be around 2.3 ms. As a comparison, the mean detection latency (determined by our simulation) with the same parameters and eye tracking system, but tracking at 1000 Hz, amounted to 3.4 ms (SD = 0.66 ms), while remaining at a high accuracy level (p(FA) = 0.3%, SD = 0.06%; mean d’ = 6.1, SD = 0.4). The second crucial latency for gaze-contingent displays is the flip latency (i.e., the time the flip occurs relative to saccade onset, see Figure 7b), which depends on the hardware, the display frame rate used, as well as the time within the refresh cycle. In this experiment, this flip latency was on average 11 ms (SD = 3.1 ms), as a mean detect-to-flip delay of 6 ms (SD = 2.46 ms) incurred. Finally, the flip-to-display latency should be deterministic, as the ProPixx DLP projector updates its entire display with microsecond precision as soon as all video data is transferred. Indeed, the flip-to-display latency amounted to an average of 8.15 ms (SD = 0.35 ms), which is in line with the graphic card’s refresh rate of 120 Hz. The addition of all system delays resulted in a physical stimulus onset around 19.6 ms (SD = 3.1 ms) after saccade onset (Photodiode onset, Figure 7b), leading to the intended intra-saccadic display right when the eye was in mid-flight (Figure 7a).

**Figure 7.**
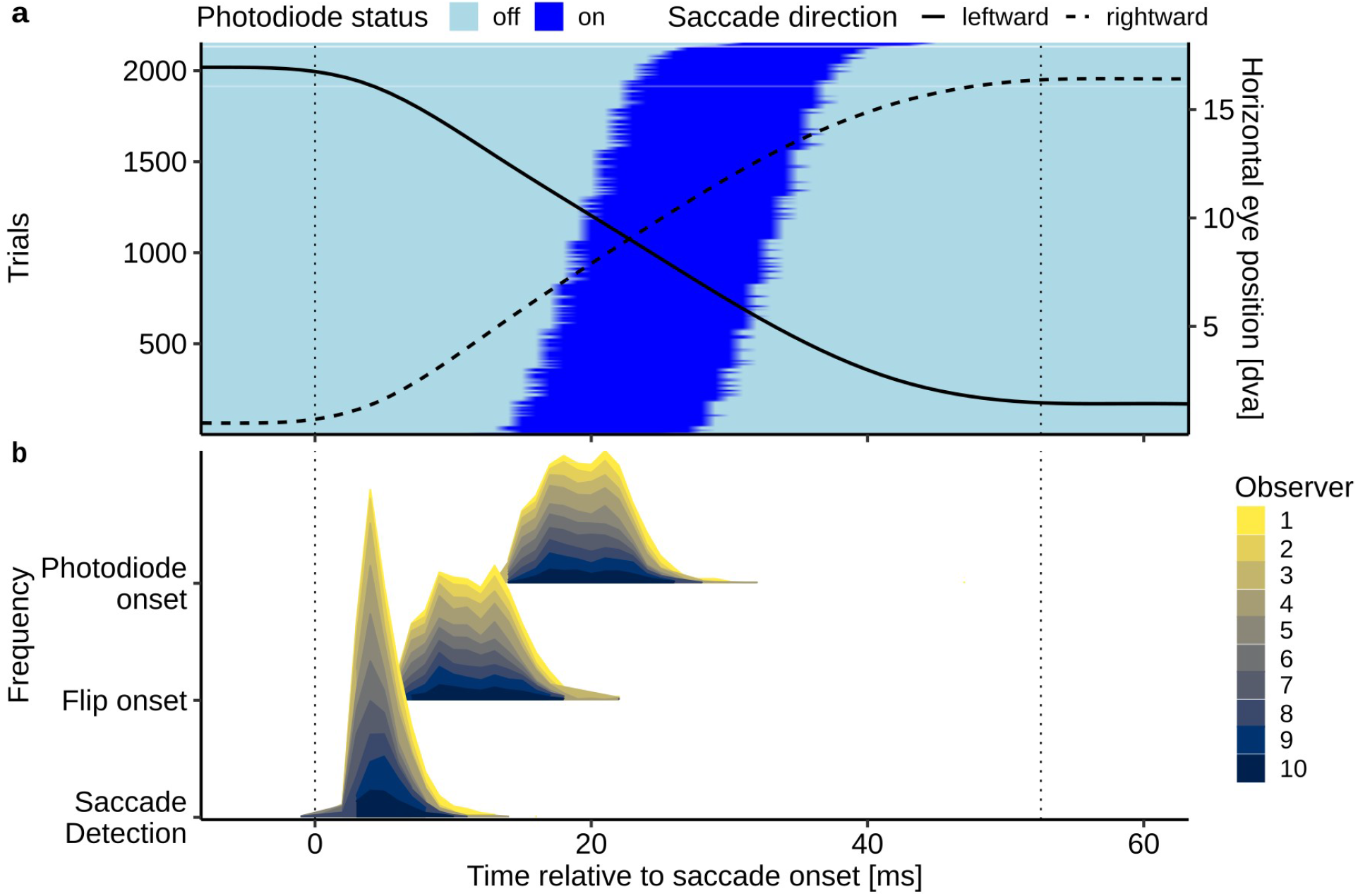
Timing event in the experimental test. **a** On-times of the photodiode (dark blue) of all saccade trials displayed and sorted relative to the onset and offset of the saccade (dotted vertical lines). Solid and dashed lines represent the mean horizontal saccade trajectories of leftward and rightward saccades over time (smoothed by a univariate general additive model). **b** Distributions of detection, flip, and photodiode onset times relative to the onset of the saccade. Shadings indicate data from different observers.

If the intra-saccadic stimulus presentation was indeed successful, then observers should have been able to detect the rapidly drifting Gabor during saccades and not during fixation, as only an ongoing eye movement could decrease the (relative) velocity of the stimulus on the retina to the point at which it would become perceivable. When observers were fixating, their detection performance was not significantly different from chance level (d’ = 0.06, SD = 0.14; t(9) = 1.28, p = .23), suggesting that the Gabor stimulus drifting at an average velocity of 419 dva/s (SD = 61.4 dva/ s) was indeed invisible when the eye was not moving (Figure 8a). In contrast, we found that stimuli were readily detected during saccades, as performance drastically improved relative to the fixation condition (d’ = 2.94, SD = 1.1; F(1,9) = 59.6, η^2^=−0.79, p < 0.0001).

**Figure 8.**
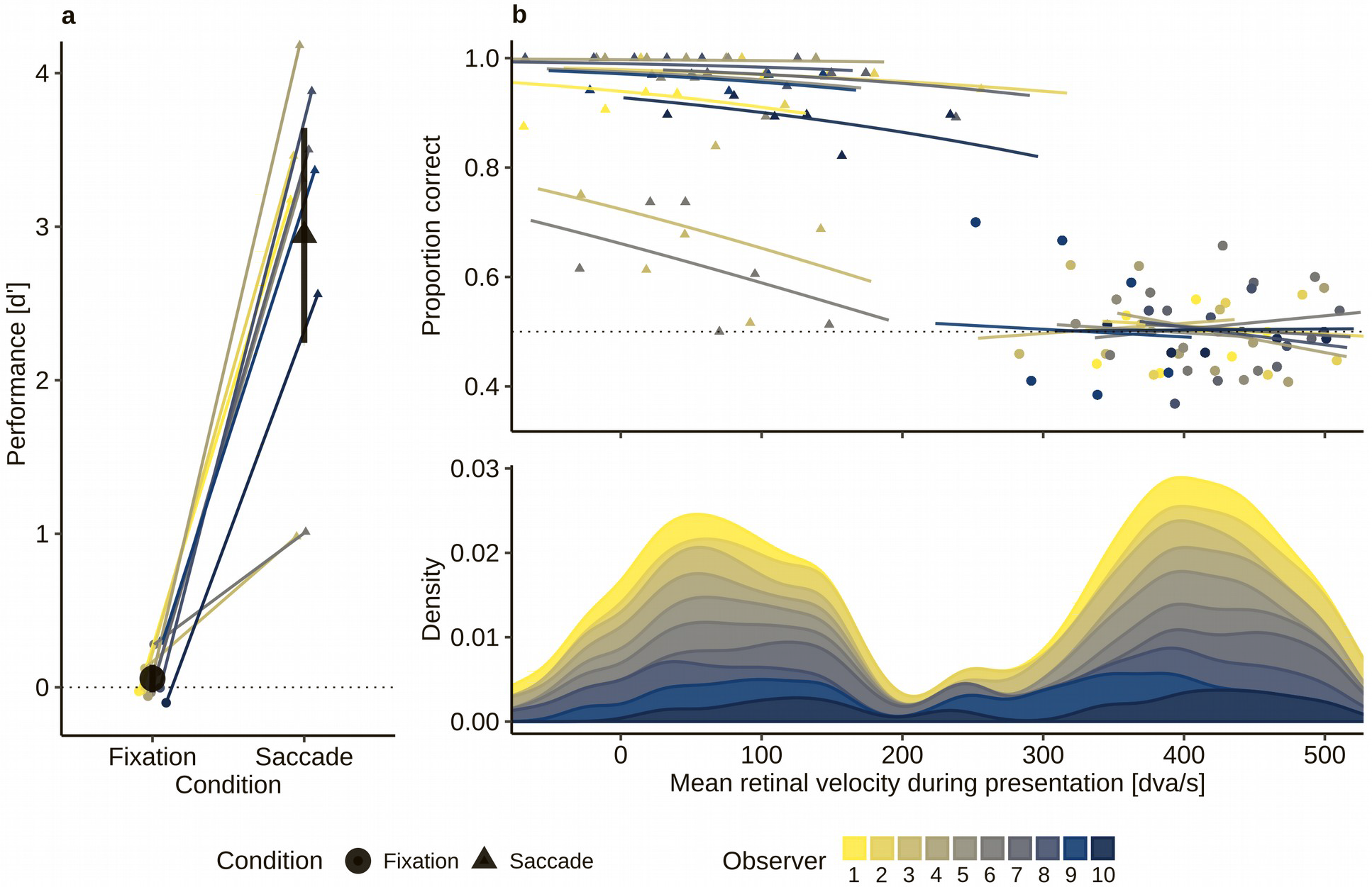
Behavioral results from the perceptual test. **a** Stimulus detection performance (d’) in fixation and saccade conditions for individual observers. Large black circles and triangles represent group means for fixation and saccade conditions. Error bars indicate 95% confidence intervals, based on ±2SEM. **b** Upper panel: Model fits from the logistic mixed-effects regression with random intercepts and slopes for observers. Points indicate mean proportion correct in six equal-sized bins of retinal velocity (i.e., mean eye velocity during stimulus presentation subtracted from stimulus velocity) per condition per observer. Lower panel: Distribution of retinal velocities for each observer for fixation (dashed lines) and saccade (solid lines) conditions.

To further explore the potential effect of retinal velocity on detection performance, we computed each trial’s mean retinal velocity during stimulus presentation (see *Analysis*). We found that retinal velocity was on average 416 dva/s (SD = 45 dva/s) during fixation, whereas it was reduced to 68 dva/s (SD = 68.8 dva/s) during saccades. Note that mean retinal velocity during saccades was in most cases positive, because the presentation of the stimulus extended into the deceleration phase of the saccadic velocity profile (Figure 7a).

A logistic mixed-effects regression (Bates, Mächler, Bolker, & Walker, 2015) revealed not only a large increase of correct responses in the saccade condition (β = 3.0, t = 5.7, 95% CI [2.13, 4.13]), but also a significant negative effect of retinal velocity (β = −0.12, t = −2.41, 95% CI [−0.23, - 0.016]), suggesting that, across both conditions, higher retinal velocity was associated with lower task performance. An interaction between condition and retinal velocity was also significant (β = −0.20, t = −2.06, 95% CI [−0.4, −0.01]). As overall performance was much lower in the fixation condition, this interaction suggests that the effect of retinal velocity was exclusive to the saccade condition (Figure 8b). To check whether the difference between fixation and saccade condition was mediated by a difference in retinal eccentricity at the time of stimulus presentation, we also computed mean retinal position of the stimulus. In the saccade condition, stimuli had an average 1 dva offset of horizontal eccentricity against the saccade direction (M_x,sac,leftward_= 1.31 dva, SD_x,sac,leftward_ = 1.1 dva; M_x,sac,rightward_ = −1.15 dva; SD_x,sac,rightward_ = 1.2 dva) relative to the fixation condition (M_x,fix,leftward_ = 0.30 dva; SD_x,fix,leftward_ = 0.54 dva; M_x,fix,rightward_ = −0.37 dva, SD_x,fix,rightward_ = 0.62 dva), which can be explained by the fact that stimulus presentation extended into the second half of the saccade in most cases, that is when the eye had already passed the screen center (the mean horizontal presentation location). To determine whether this slight difference in eccentricity had any effect on task performance, we added absolute horizontal and vertical eccentricity to the logistic mixed-effects regression. We found an increase in log-likelihood that was not significant (ΔLL=+8.8, χ^2^(21)=17.51, p=.68), suggesting that retinal eccentricity played, if any, a subordinate role in our task.

## Discussion

Timing is crucial when studying visual perception around the time of saccades, especially when manipulating stimulus configurations gaze-contingently with the onset of a saccade. In gaze-contingent experimental paradigms, various sources of latency have to be considered, ranging from eye tracker delays, saccade detection latencies, graphic card refreshes, video delays, and monitor reaction time. Whereas most of these latencies can be reduced by enhancing hardware capabilities, a more efficient online saccade detection algorithm improves timing performance independently of the experimental setup. Unfortunately, the most commonly used online detection techniques, i.e., the spatial boundary technique and the absolute velocity threshold, have significant shortcomings: The former provides reliable but late saccade detection, whereas the latter is fast but struggles with reliability, especially at higher sampling rates or slightly increased noise levels. Inspired by a widely-used algorithm for (micro-)saccade detection (Engbert & Kliegl, 2003; Engbert & Mergenthaler, 2006), we developed a velocity-based online saccade detection algorithm that incorporates both algorithms’ strong points: It allows for rapid saccade detection due to low velocity thresholds, it is robust against noise by applying smoothing, adaptive adjustment of velocity thresholds, and an optional direction criterion, and it allows the user to flexibly specify more liberal or more conservative detection criteria. Across various gaze-contingent experimental paradigms, as well as in non-scientific applications, this open-source algorithm could help create comparable and reproducible results by (1) avoiding timing problems (due to its early saccade detection) and (2) increasing stability (due to its increased robustness against noise).

We validated the algorithm, as well as the boundary and the absolute-velocity technique, in a large-scale simulation (>30,000 saccades). We found that the algorithm provided considerably earlier saccade detection than boundary techniques (up to 10 ms or more, depending on sampling rate), which was more similar to (although in most cases slightly slower than) the latency of the absolute velocity technique. Crucially, the algorithm’s accuracy in online saccade detection was on par with the boundary technique and significantly larger than that of the absolute velocity technique, especially at high sampling rates. Moreover, when corrupting the collected data with noise, absolute velocity techniques suffered from a drastic increase in false alarms, whereas the proposed algorithm maintained its detection accuracy by updating its velocity threshold. This is an important result and prerequisite for the use during eye tracking experiments, because many factors that vary throughout the experiment or on a trial-by-trial basis, such as pupil diameter, time since last calibration, head movements, gaze eccentricity, or marker visibility, may influence accuracy and precision of recordings (Nyström, Andersson, Holmqvist, & Van De Weijer, 2013) and thus alter the noise level. Since the algorithm updates velocity thresholds with every new incoming data sample, various negative influences on detection accuracy can be accounted for. At the same time, the incentive to achieve high data quality remains, as increased noise levels may still have a negative impact on detection latency. Running the algorithm in real-time is feasible as its implementation can process a large number of collected samples, such as those collected from eye trackers sampling at 2000 Hz, within microseconds. However, due to its linear time complexity, it cannot be guaranteed that the algorithm finishes prior to any given deadline. For example, if the algorithm were run on an unnecessary large number of samples, such as 4,000,000, then it would take ~187 ms (based on the results of our simulation), exceeding the duration of a saccade by far.

We found that the detection criterion and subsequent performance of the algorithm strongly depends on the parameters supplied: More conservative settings (i.e., higher threshold factor, more samples above threshold, a tighter range of accepted directions) will improve detection accuracy to a maximum, but will come at the cost of increased detection latency, and in the worst case—as shown by the high noise conditions—may lead to abnormally high velocity thresholds that will make saccade detection impossible. We thus suggest a careful weighting of parameters depending on the experimental setup and paradigm used. For example, when using a threshold factor of λ=5, it makes sense to have at least three samples above threshold to detect a saccade. Indeed, if the algorithm were used to detect microsaccades during an experiment, low thresholds should be used as eye velocities during microsaccades are not as high as during saccades, while three or more samples should be evaluated (for a successful application with a different implementation, see Yuval-Greenberg, Merriam, & Heeger, 2014). If however a threshold factor of λ=20 is used, one sample above threshold may often be enough to reliably detect a saccade without adding significant additional delay. In pilot work with the TrackPixx3 (VPixx Technologies, 2017), we also found that binocular online saccade detection (running the detection algorithm on each eye separately) allows for lower threshold factors, as the probability that velocity thresholds are exceeded due to noise in both eyes simultaneously is smaller than for one eye only. The choice of threshold also depends on the noise level and sampling rate of the eye tracker in use, as a higher sampling rate can inflate velocity estimates (Han, Saunders, Woods, & Luo, 2013). In some systems, such as the Eyelink 1000+ (SR-Research, 2013), these two variables are not independent: With deactivated heuristic filters, root-mean-square noise amounts to 0.02 dva at 1000 Hz and to 0.03 dva (monocular tracking) and 0.04 dva (binocular tracking) at 2000 Hz. Additional filter levels supplied by the manufacturer can reduce these noise levels, but introduce additional end-to-end sample delay (also depending on sampling rate). It is thus important to understand that the parameter choice for optimal online saccade detection performance is intrinsically dependent on both the recording settings and the nature of the task. For instance, a direction restriction can only be applied in paradigms, in which it is certain or at least very likely that the participant will indeed make a saccade in a given direction. In case of a two-alternative forced choice paradigm, it would be possible to call the algorithm twice, each time with different direction criteria. But it would be impossible to apply a direction criterion in a free viewing context. Ultimately, it remains an advantage that the experimenter is able to fine-tune the detection criterion according to the relative costs for longer detection latencies or for an increased false alarm rate incurring in a specific task.

With the introduction of the objective and perceptual test experiment, we provided a real-world example of a gaze-contingent paradigm in which timing was crucial, in this case for the intra-saccadic presentation to be successful. This test builds on the finding that rapidly drifting or flickering gratings that are invisible during fixation can be rendered visible due to the reduction of retinal velocity occurring when the eye moves across them (Castet & Masson, 2000; Mathôt, Melmi, & Castet, 2015; Garcìa-Pérez & Peli, 2011). We used a projection system operating at submillisecond temporal resolution to briefly display a rapidly-drifting Gabor patch entirely during the saccade, and we asked observers to detect it. We established that the Gabor patch was indeed largely invisible during fixation. It was absolutely crucial in this task, therefore, that the stimulus was presented while the eye was in mid-flight to achieve an approximate match of the velocities of stimulus and eye. Both observers’ high task performance in the saccade condition (perceptual test) and recordings from a photodiode (objective test) confirmed that despite all possible system delays strictly intra-saccadic presentations with a physical onset as early as 20 ms after saccade onset were well possible. At the same time, only 1% of all trials had to be removed from analysis because presentations did not happen strictly during the saccade, e.g., due to late detections or erroneous detections while fixating (see Methods).

But the finding is interesting for two other reasons. First, it shows that if a stimulus has high contrast, is optimized for the high velocity of saccades (Castet & Masson, 2000; Deubel, Elsner, & Hauske, 1987; Garcìa-Pérez & Peli, 2011; Mathôt, Melmi, & Castet, 2015; Watson, Schweitzer, Castet, Ohl, & Rolfs, 2017), and is not affected by pre- and post-saccadic masking (Campbell & Wurtz, 1978; Castet, 2010), then it is readily detectable, if not highly salient. Second, the finding indicates that timing during and around saccades matters. It is widely assumed that visual processing is suppressed during and around the time of saccades (Burr, Holt, Johnstone, & Ross, 1982; Burr, Morrone, & Ross, 1994; Ross, Morrone, Goldberg, & Burr, 2001). This is the reason why many trans-saccadic paradigms relying on gaze-contingent manipulations assume that as long as a display change that falls within the window of saccadic suppression—which precedes saccade onset by up to 100 ms and exceeds saccade duration by up to 50 ms (Diamond, Ross, & Morrone, 2000; Volkmann, Riggs, White, & Moore, 1978; Volkmann, 1986)—is neither noticed nor processed. There is, however, converging evidence that stimuli undergoing saccadic suppression can shape post-saccadic perception (Watson & Krekelberg, 2009), and that the relative timing of a stimulus relative to saccade offset drastically changes both the appearance of that stimulus and its likelihood to be consciously perceived (Balsdon, Schweitzer, Watson, & Rolfs, 2018; Bedell & Yang, 2001; Campbell & Wurtz, 1978; Duyck, Collins, & Wexler, 2016; Matin, Clymer, & Matin, 1972). Although for many experiments and the conclusions drawn from them intra-saccadic display changes may not be absolutely crucial, it is important to be aware of the possibility that an intended intra-saccadic change might in fact be a non-intended post-saccadic change due to insufficient control of timing or hidden latencies in the hardware. Earlier saccade detection can alleviate this risk.

We conclude that implementing efficient gaze-contingent display changes across saccades can be tricky owing to a range of system latencies that have an impact on a paradigm’s timing behavior. We as experimenters need to examine these latencies closely to draw the right conclusions from our results. Early online saccade detection can assist greatly in this task, as it saves valuable time for the setup to perform the intended (trans-saccadic) changes, but it comes at the cost of reduced online saccade detection accuracy—especially at higher noise levels—making it ultimately harder to smoothly collect data. The algorithm proposed here outperforms traditional detection methods in speed and accuracy, while adjusting detection thresholds in response to increased noise levels. These properties make it a reliable tool even when collecting data even under sub-optimal recording circumstances, while being computationally feasible for the online scenario due to its near real-time processing and linear complexity. Finally, the open-source availability of the code leaves it open for every researcher to use it and adapt it to specific needs, making it a versatile tool for the field of active vision.

## Acknowledgments

We acknowledge the significant contribution of Ralf Engbert, Konstantin Mergenthaler, Petra Sinn and Hans Trukenbrod for making the code of their microsaccade detection toolbox publicly available, as well as Nicolas Devillard for the excellent ANSI C implementations and comparisons of different median search algorithms (http://ndevilla.free.fr/median/median/index.html). RS was supported by the Studienstiftung des deutschen Volkes and the Berlin School of Mind and Brain. MR was supported by the Deutsche Forschungsgemeinschaft (DFG, grants RO3579/2-1, RO3579/8-1 and RO3579/10-1).

## Author Contributions

RS implemented the algorithm and ran simulations. Validation procedure was conceptualized by RS and MR. The experimental test was designed, run, and analyzed by RS under MR’s supervision. RS drafted the manuscript and MR provided critical revisions.

## Open Practices Statement

Saccade data and code used for simulations, data collected throughout the experimental test, experimental code, as well as data analysis scripts are available on the Open Science Framework: https://osf.io/3pck5/. The experimental test was not preregistered.

Implementations of the proposed algorithm in C, Python, and Matlab are available on Github: https://github.com/richardschweitzer/OnlineSaccadeDetection.

